# Normalized coefficient of variation (nCV): a method to evaluate circadian clock robustness in population scale data

**DOI:** 10.1101/2021.07.28.454045

**Authors:** Gang Wu, Lauren J. Francey, Marc D. Ruben, John B. Hogenesch

## Abstract

Robust oscillation of clock genes is a core feature of the circadian system. Relative amplitude (rAMP) measures the robustness of clock gene oscillations, but only works for longitudinal samples. We lack a method for estimating robust oscillations from human samples without labeled time. We show that the normalized coefficient of variation (nCV) is linearly correlated with rAMP, independent of time labels. Using nCV, we found that clock gene oscillations are consistently dampened in tumors compared to non-tumors, suggesting a new therapeutic target in cancer treatment by enhancing clock robustness. nCV can provide a simple measure of the robustness of clock gene oscillations in any population-level dataset.

**Availability and implementation:** The nCV web application is available on the GitHub repository (https://github.com/gangwug/nCV).

## Background

The negative feedback loop of the clock gene network drives robust clock gene oscillations (Gonze *et al.*, 2002; Ukai and Ueda, 2010). The robustness of clock gene oscillations (hereafter referred to as clock robustness) can be thought of as relative amplitude (rAMP, the magnitude of the wave). Clock gene knockouts (e.g., *Arntl, Per1/2,* and *Cry1/2*) and disease states (e.g., cancers, metabolic syndrome and sleep disorders) can reduce clock robustness (Bunger *et al.*, 2000; Bae *et al.*, 2001; Kume *et al.*, 1999; Anafi *et al.*, 2017). Therefore, it is important to measure clock robustness. The rAMP calculation requires time labeled samples (Wu, Ruben, Lee, *et al.*, 2020). However, most human samples lack time information. There are currently no methods to directly measure clock robustness in human samples when the time of collection is unknown.

We show that the normalized coefficient of variation (nCV) is linearly correlated with rAMP for clock genes in longitudinal mouse and human datasets. Importantly, nCV can be applied to *unordered* datasets. Therefore, we applied nCV to analyze multiple cancer types and found that clock robustness is consistently reduced in tumors compared to adjacent non-tumor samples.

## Results and Discussion

Many lines of evidence suggest that the clock is dysregulated in cancer (Sancar and Van Gelder, 2021). rAMP indicates more robust circadian oscillation of *ARNTL* in normal human breast epithelial cells (MCF10A; rAMP = 0.085) compared to breast cancer cells (MCF7; rAMP = 0.003) (Fig. 1A). The rAMP calculation requires the sample collection time. Other metrics, such as the clock correlation distance (Shilts *et al.*, 2018), can indicate clock function in the absence of collection time, but cannot measure clock robustness (Fig. S1). The majority of patient datasets available do not have recorded collection times. Therefore, we need a way to estimate clock robustness in population data where sampling time is not recorded.

**Fig. 1.**
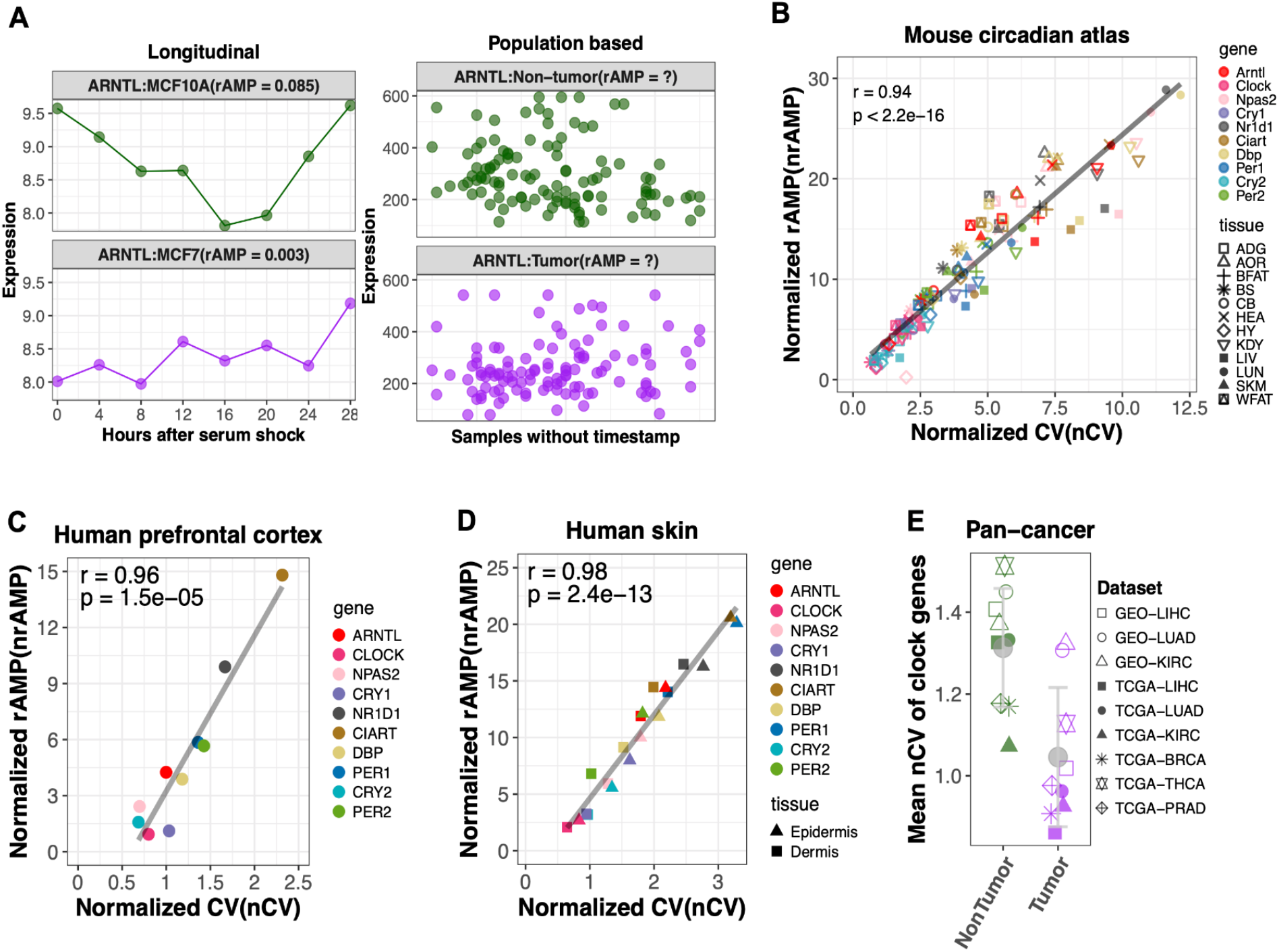
nCV measures clock robustness. (A) *ARNTL* oscillations are stronger (higher rAMP)) in breast epithelial cells (MCF10A; green) than breast cancer cells (MCF7; purple). rAMP is uninformative when comparing human breast tumors versus adjacent non-tumor samples without labeled time (right panel). (B) nCV of clock genes is linearly correlated with nrAMP in 12 mouse tissues. (C) nCV is linearly correlated with nrAMP in the human prefrontal cortex samples with labeled time. (D) nCV is linearly correlated with nrAMP in the human epidermis and dermis population samples without labeled time but ordered by CYCLOPS. (E) Reduced clock robustness in tumor versus adjacent non-tumor tissues. The mean nCV of 10 clock genes from tumor and non-tumor samples for each dataset is indicated by green and purple points respectively. Grey point indicates the average value (+/SD) in non-tumor vs. tumor group.

To address this, we selected a group of 10 clock genes that: 1) are important in circadian regulation, 2) cycle in multiple tissues, 3) are phased across the full circadian cycle, and 4) represent a range of rAMP values (Fig. S2). Then, we searched for a time-independent measure of variance that is linearly correlated with rAMP. Median absolute deviation (MAD), standard deviation (SD), and the coefficient of variation (CV) are common measures of transcript level variance. We found that the CV correlates with rAMP (r = 0.96) much better than with either MAD (r = 0.35) or SD (r = 0.37) for circadian genes in wild type mouse liver (Fig. S3). We used the normalized CV (nCV), the ratio between the CV of a clock gene and the mean CV of all genes to adjust for systemic difference in CV between datasets. To validate this new metric, we compared nCV to the normalized rAMP (nrAMP) for a variety of datasets.

The nCVs of the 10 clock genes show strong linear correlation (r = 0.94 and p < 2.2e-16) to nrAMPs in 12 mouse tissues (Fig. 1B). Importantly, the nCV is reduced in several genetic knockout models of the clock (*Cry1*/*Cry2* double knockout mouse livers; *Arntl* knockout cells and mouse tissues; *Clock* knockout mouse kidney; *Nr1d1*/*Nr1d2* double knockout mouse livers) (Fig. S4). The mean nCV of 10 clock genes in the knockout group is smaller (p = 2.3e-6) than the wild type group (Fig. S5). We next validated nCV in human samples with labeled time. nCV is strongly correlated with nrAMPs in human prefrontal cortex samples (Fig. 1C; r = 0.96 and p = 1.5e-05) (Chen *et al.*, 2016) and human skin samples ordered by CYCLOPS (Fig. 1D; r = 0.98 and p = 2.4e-13) (Wu, Ruben, Francey, *et al.*, 2020). In sum, nCV is a strong surrogate measure for nrAMP even when sample collection time is unknown.

The vast majority of human datasets do not have time of day information available. This includes most cancer datasets. Thus, we applied nCV to study clock robustness in tumor versus adjacent non-tumor samples without labeled time. Clock robustness was reduced in tumor samples from patients with hepatocellular carcinoma, lung adenocarcinoma, kidney renal clear cell carcinoma, breast cancer, and thyroid carcinoma (Fig. S6). *ARNTL* and *PER2* oscillations are consistently decreased in tumor samples from all 9 tested datasets. In sum, clock robustness is reduced (p = 0.002) in tumors compared to non-tumors in all 9 datasets (Fig. 1E).

Studies report inconsistent changes in clock gene expression between different human cancer types (Savvidis and Koutsilieris, 2012; Ye *et al.*, 2018). However, we show that clock robustness using nCV is consistently reduced across human cancers. Clock robustness may in fact be an important therapeutic target in cancer. For example, robust oscillation of *Per2* in SCN is essential to drive locomotor activity rhythms (Chen *et al.*, 2009). In addition, enhancing the circadian clock function in cancer cells inhibits tumor growth and improves cardiometabolic health for patients with metabolic syndrome (Kiessling *et al.*, 2017; Wilkinson *et al.*, 2020). Beyond cancer, nCV can be used to estimate clock robustness in any context where population-scale data are available. In sum, nCV provides a straightforward measure of circadian clock function in the absence of labeled time, the vast majority of all human datasets.

## Methods

All datasets used in this study are available in the GEO database and Firebrowse (Table S1). All statistical analyses were performed in R.

## Acknowledgments

We thank Drs. Andrew Liu, Eric Zhang, and Rafael Irizarry for helpful discussions about circadian amplitude. We thank Ron Anafi for discussing these issues.

