## Supplemental file 1 for "Normalized coefficient of variation (nCV): a method to evaluate circadian clock robustness in population scale data"

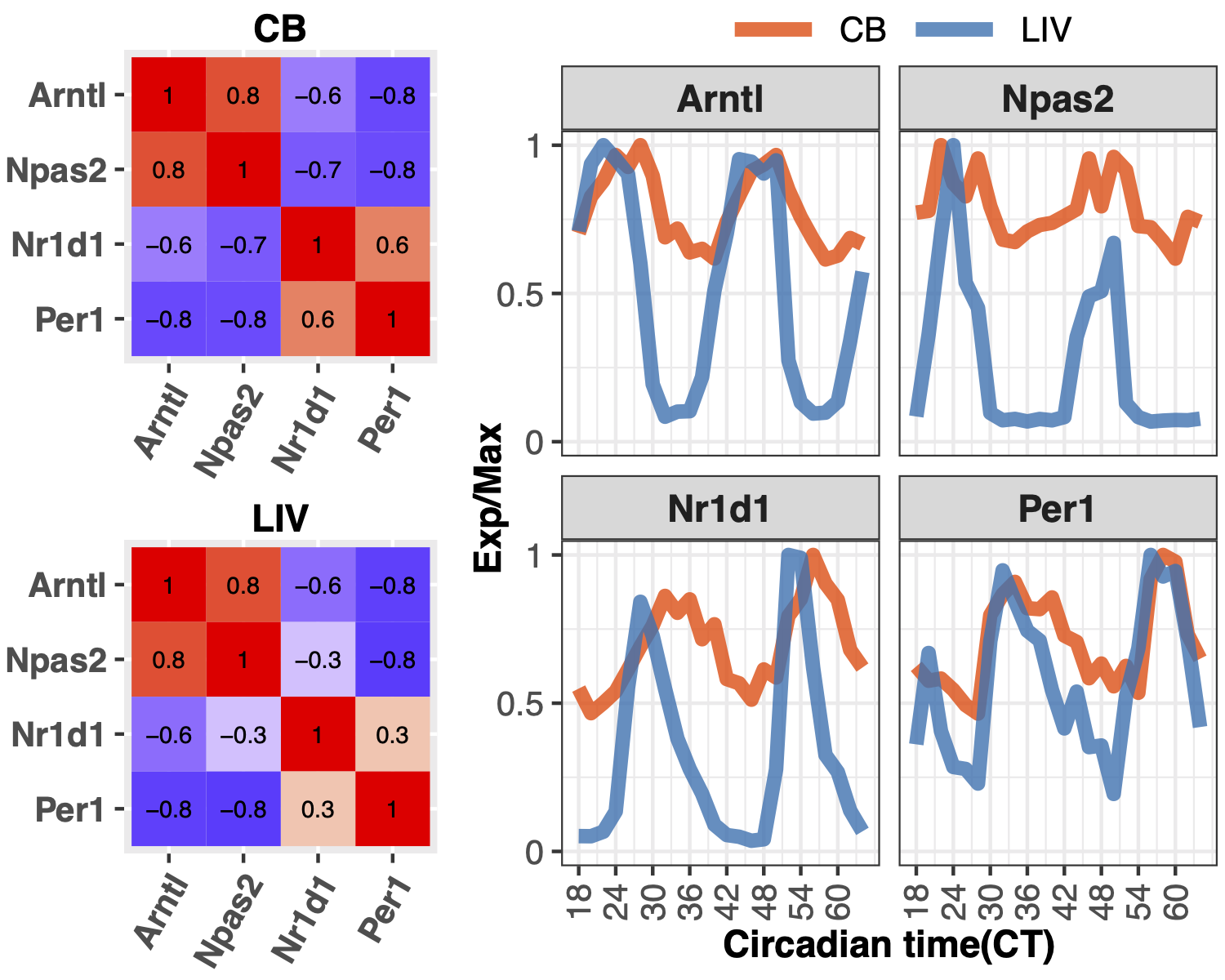

**Fig. S1.** Spearman’s ρ does not indicate the robustness of clock gene oscillation. Left panel, Spearman’s ρ for *Arntl*, *Npas2*, *Nr1d1* and *Per1* from time-series data of mouse cerebellum (CB) and liver (LIV). Red and blue color indicate positive and negative Spearman’s ρ for each pair of clock genes. Right panel, time-series expression profiles from mouse cerebellum (orange) and liver (blue). Exp/Max indicates the expression value at each time point normalized to the maximum expression across time points.

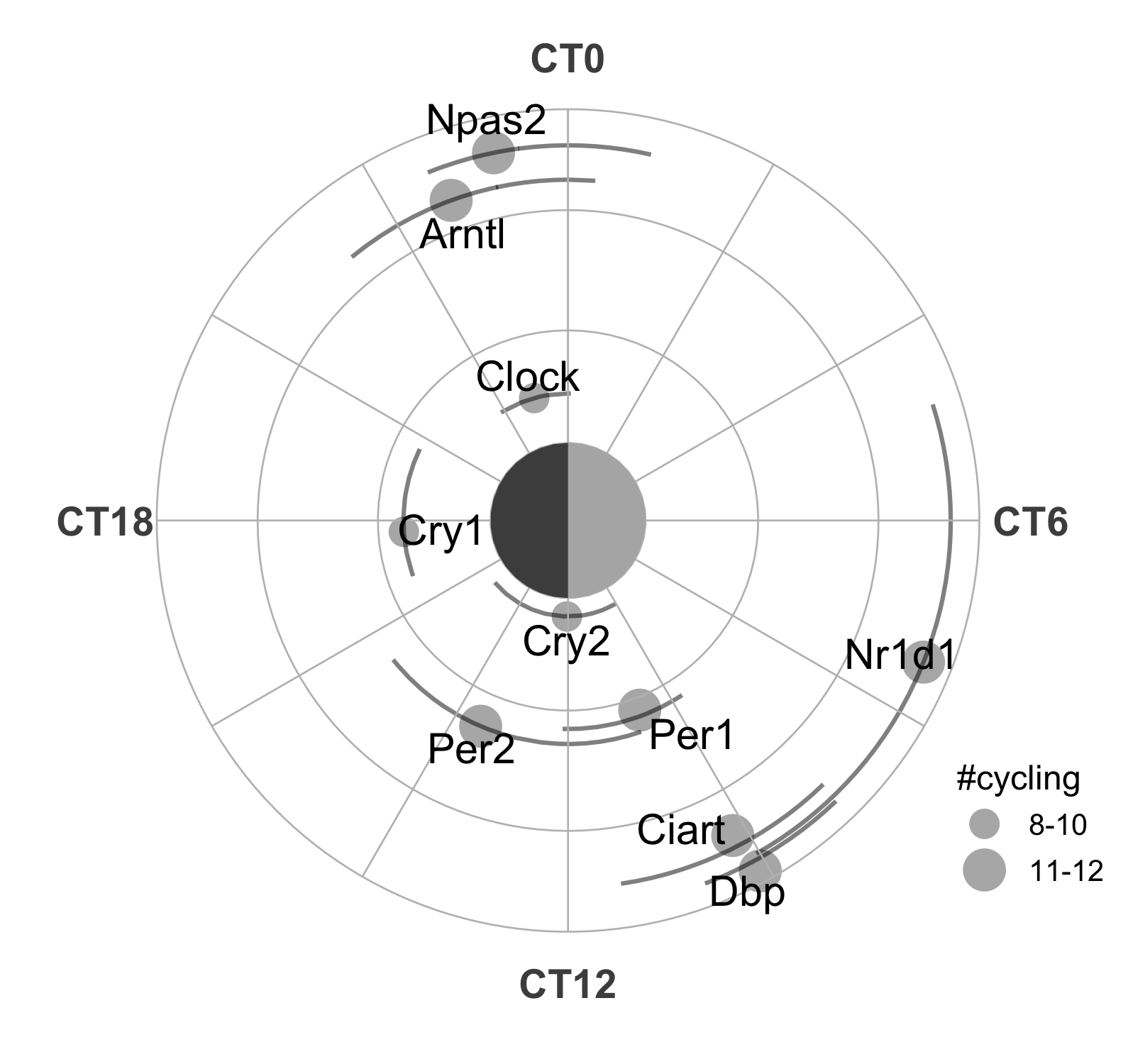

**Fig. S2.** The phase and rAMP of clock genes in 12 mouse tissues. Point size indicates the number of tissues that a clock gene is cycling (BHQ < 0.05, rAMP > 0.1). The average phase and rAMP of each clock gene among tissues are indicated by the angle and the radius length. The curve length indicates the phase range of a clock spreading among tissues.

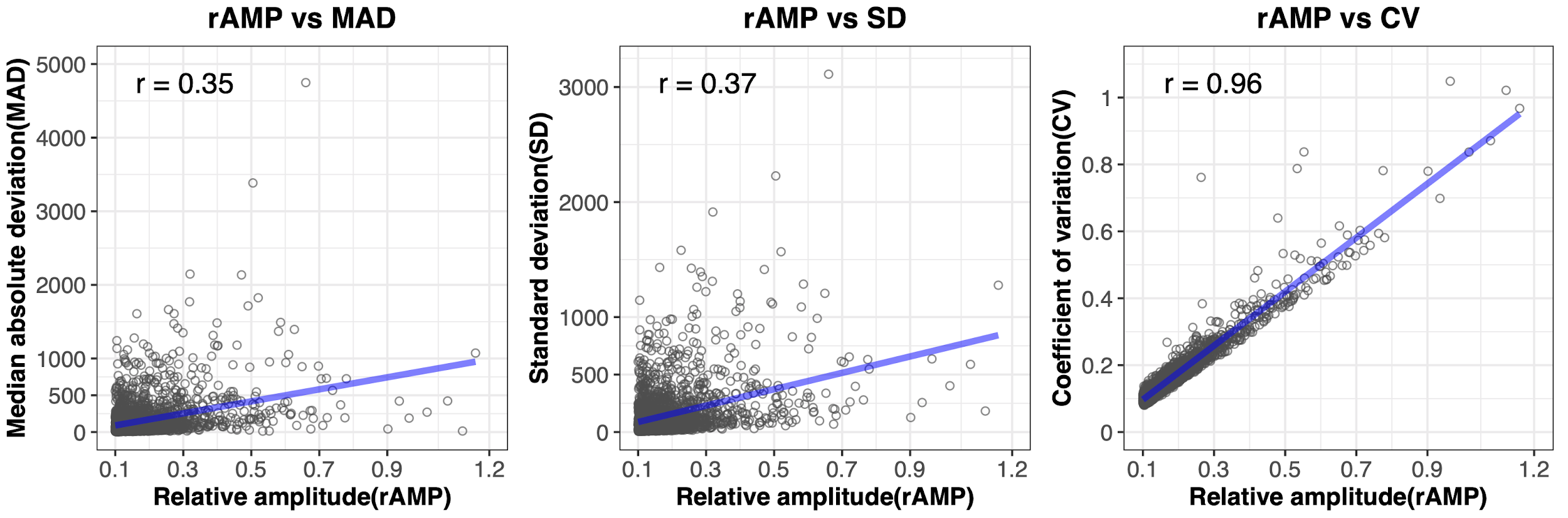

**Fig. S3.** rAMP correlates far better with coefficient of variation (CV) than median absolute deviation (MAD) or standard deviation (SD). Each point is a cycling gene (BHQ < 0.05, rAMP > 0.1) in the mouse liver identified by MetaCycle.

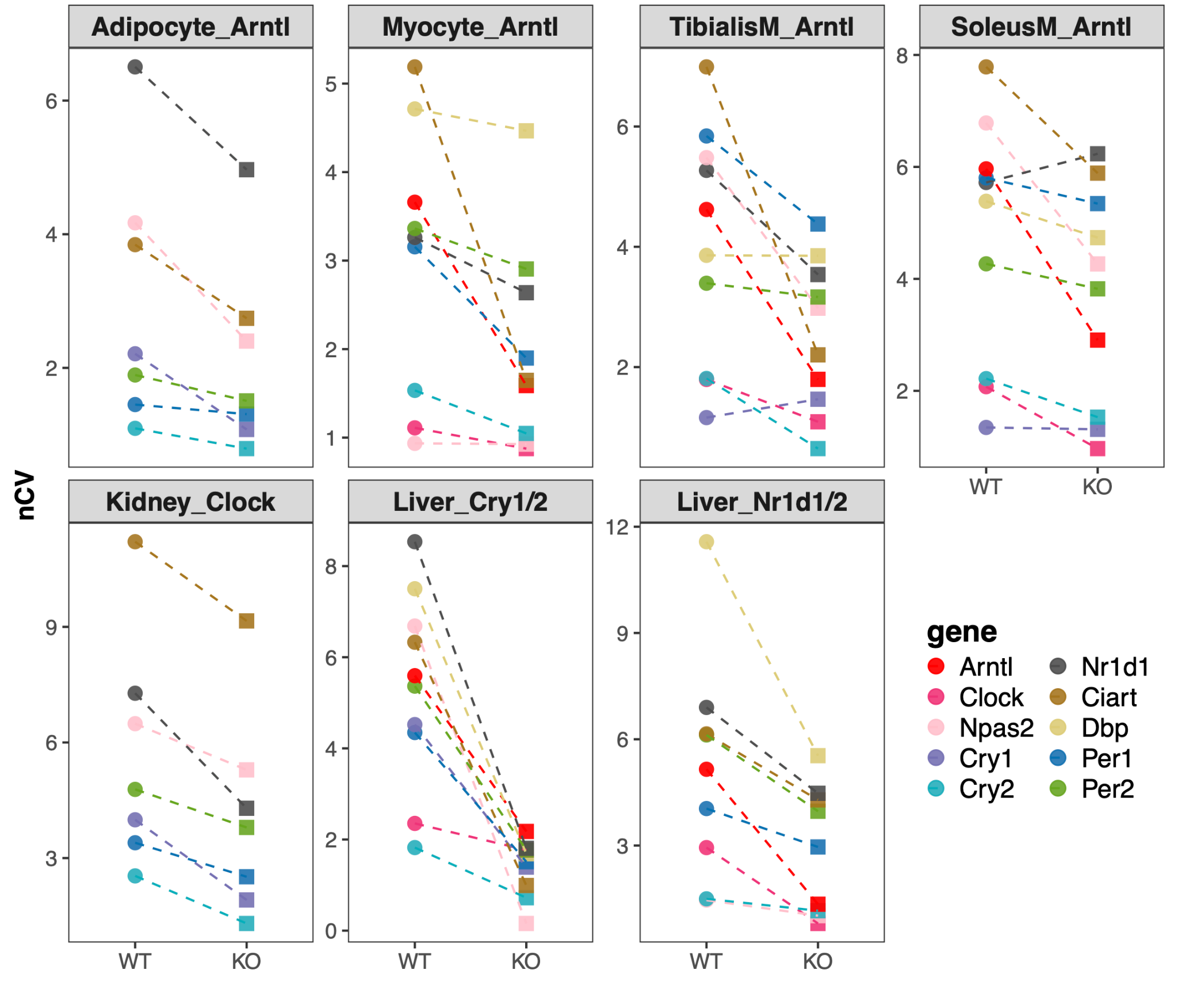

**Fig. S4.** Reduced nCV of clock genes in multiple tissues/cells with clock gene knockout compared to wild type. Clock genes are indicated by point color. The datasets include wildtype (WT) and *Arntl* knockout (KO) adipocytes (Adipocyte_Arntl), cardiomyocyte (Myocyte_Arntl), Tibiallas muscle (TibiallasM_Arntl) and Soleus muscle (SoleusM_Arntl), wildtype and *Clock* knockout kidney (Kidney_Clock), wildtype and *Cry1*/*Cry2* double knockout liver (Liver_Cry1/2), and wildtype and *Nr1d1*/*Nr1d2* double knockout liver (Liver_Nr1d1/2). The mean nCV of 10 clock genes of wild type and knock-out cell/tissues is indicated by green and purple points respectively.

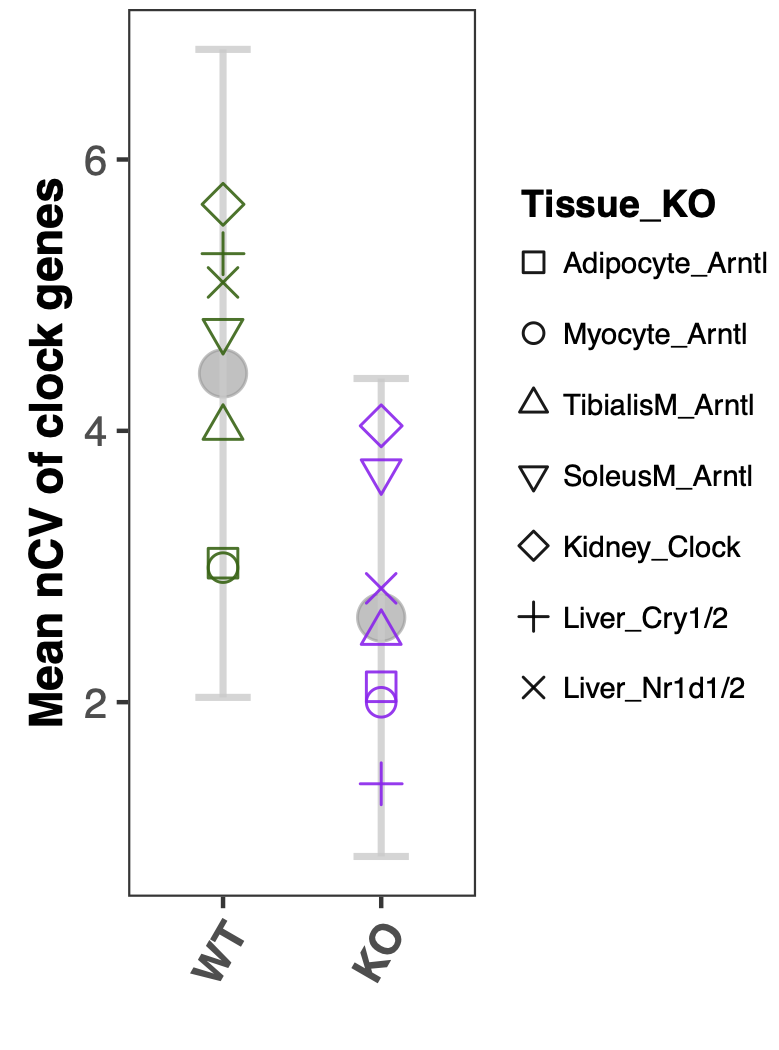

**Fig. S5.** nCV reports reduced robustness of circadian clock in clock gene mutant cell/tissues compared to wild type cell/tissues. Point shape indicates the dataset, including wildtype and *Arntl* knock-out adipocytes (Adipocyte_Arntl), cardiomyocyte (Myocyte_Arntl), Tibiallas muscle (TibiallasM_Arntl) and Soleus muscle (SoleusM_Arntl), wildtype and *Clock* knock-out kidney (Kidney_Clock), wildtype and *Cry1*/*Cry2* double knock-out liver (Liver_Cry1/2), and wildtype and *Nr1d1*/*Nr1d2* double knockout liver (Liver_Nr1d1/2). The mean nCV of 10 clock genes of wild type and knock-out cell/tissues is indicated by green and purple points respectively. Grey point indicates the average value in wildtype or knockout group, with error bar indicating +/- SD.

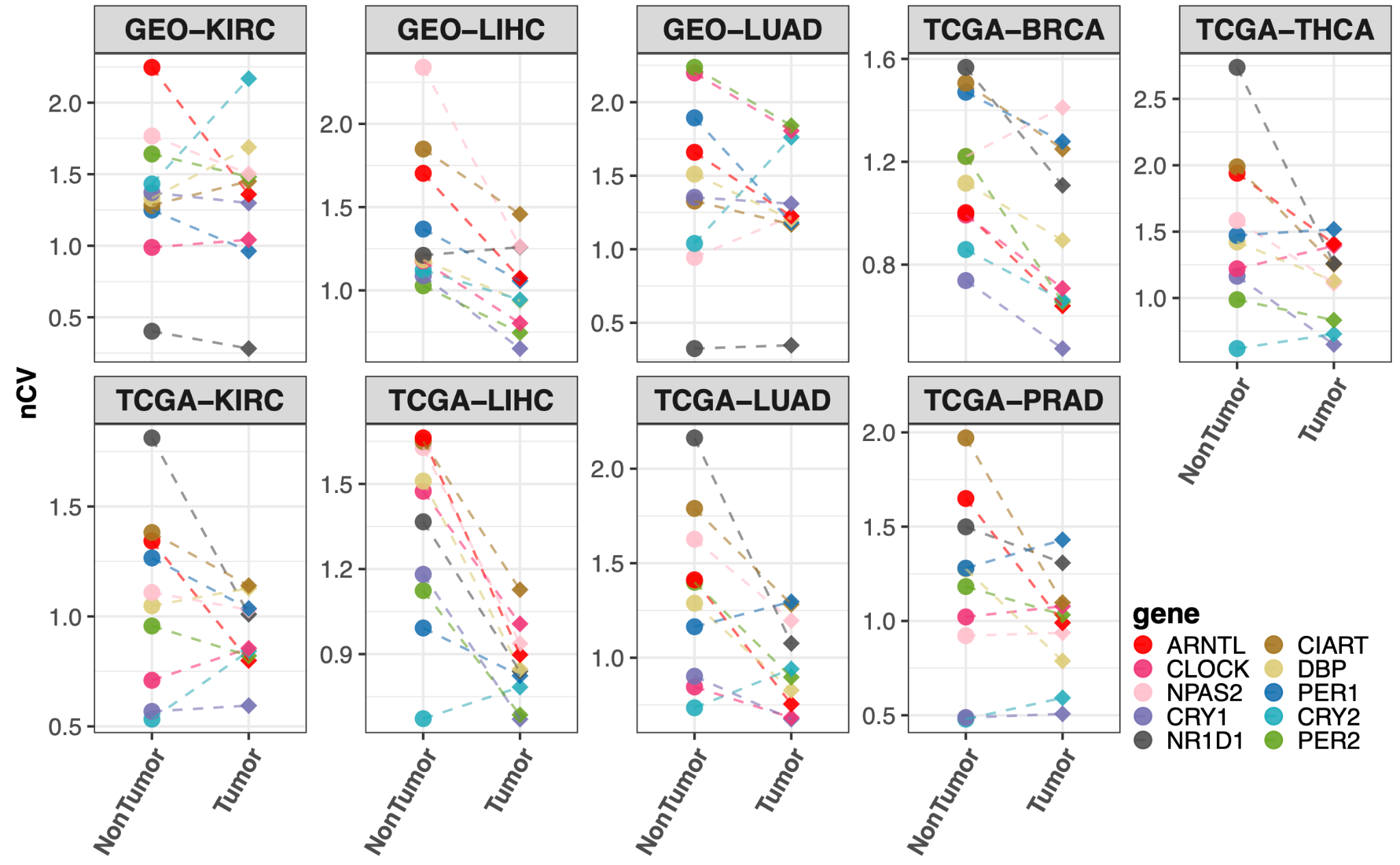

**Fig. S6.** nCV of 10 clock genes in tumor and adjacent non-tumor samples in datasets from GEO and TCGA database. Reduced nCV of clock genes in kidney (GEO-KIRC), liver (GEO-LIHC) and lung (GEO-LUAD) from GEO database, breast (TCGA-BRCA), thyroid (TCGA-THCA), kidney (TCGA-KIRC), liver (TCGA-LIHC), lung (TCGA-LUAD) and prostate adenocarcinoma (TCGA-PRAD) tumor compared to adjacent non-tumor samples from TCGA database. The clock genes are indicated by point color. The clock genes are indicated by point color.

**Table S1: The list of datasets used in this study**

| **Datasets** | **Reference** | **Experimental design** |
| --- | --- | --- |
| 1. Mouse circadian atlas  (Longitudinal) | Zhang R, et al., 2014, PNAS. GSE54650 | 288 samples total covering 12 different tissues, with no replicates. Each tissue sampled every 2 hours for 2 days (24 samples per tissue). |
| 2. Mouse Adipocyte_Arntl (Longitudinal) | Paschos GK, et al., 2012, Nat Med.  GSE35026 | Adipose tissues were isolated from inguinal adipose fat pads at four different times of the diurnal cycle under constant darkness (circadian time, CT) for RNA extraction and hybridization on Affymetrix microarrays. Adipocyte-specific Bmal1 KO (Ad-Bmal1-/-) and control mice were used for isolation of tissues at CT0, CT6, CT12, CT18 where CT0 the beginning of the subjective day. |
| 3. Mouse Myocyte_Arntl (Longitudinal) | GSE43073 | RNA from whole hearts collected every 3 hours for 24 hours from wildtype and CBK mice was isolated and analyzed using MouseRef-8_V2 BeadChips (Illumina, Inc.). The 24-hour data were examined for rhythmicity using cosinor analysis and differences in rhythmicity between genotype groups were further examined for differences in the model fitting parameters. |
| 4. Mouse TibialisM_Arntl (Longitudinal) | Dyar KA, et al., 2014, Mol Metab. GSE43071 | 72 samples were analyzed, comprising 4 experimental groups (Ctrl SOL, mKO SOL, Ctrl TA, mKO TA), with 3 biological replicates for each time point sampled every 4 hours for 24 hours. SOL and TA muscles were collected from the same animals, as indicated by Source Animal ID data column |
| 5. Mouse SoleusM_Arntl (Longitudinal) | Dyar KA, et al., 2014, Mol Metab. GSE43071 | 72 samples were analyzed, comprising 4 experimental groups (Ctrl SOL, mKO SOL, Ctrl TA, mKO TA), with 3 biological replicates for each time point sampled every 4 hours for 24 hours. SOL and TA muscles were collected from the same animals, as indicated by Source Animal ID data column |
| 6. Mouse Kidney_Clock (Longitudinal) | GSE27366 | Examine the temporal profiles of gene expression in the mouse whole kidney. Animals were sacrificed for microdissection every 4 hours, i.e. at ZT0, ZT4, ZT8, ZT12, ZT16 and ZT20 (ZT – Zeitgeber time, indicates time of light-on as ZT0 and time of light-off as ZT12). The microarray hybridization was performed in duplicates on pools of RNA composed of equivalent amounts of RNA prepared from teo or three animals at each ZT time-point. |
| 7. Mouse Liver_Cry1/2 (Longitudinal) | Vollmers C, et al., 2009, PNAS. GSE13062 | Cry1, Cry2 double KO mice were entrained either to ad libitum or temporally restricted feeding (tRF) schedules. Food was made available to mice under the tRF regimen only between ZT(CT)1 and ZT(CT)9. Mice were then released into constant darkness while the respective feeding schedules were still maintained. Liver tissue was collected on the second day of constant darkness at the indicated time points. Total RNA was extracted and 5ug of RNA was used in the standard Affymetrix protocol for amplification, labeling and hybridization |
| 8. Mouse Liver_Nr1d1/2 (Longitudinal) | Cho H, et al., 2012, Nature.  GSE34018 | Total RNA was obtained from livers of wild-type and Liver-specific Reverb alpha/beta double knockout mice at ZT 0, 4, 8, 12, 16, and 20 |
| 9. Human epidermal and dermal  samples  (longitudinal) | Wu G., et al., 2020, Genome Medicine.  GSE112660 and  GSE139300 | Four forearm skin samples were collected at 12am, 6am, 6pm, 12pm for each of 20 male participants, except one missing sample from participant 115. The ages of these 20 participants are between 21 and 49 years old. LCM was performed to separate dermis from epidermis. |
| 10. Human prefrontal cortex (longitudinal) | Chen CY et al., 2016, PNAS. GSE71620 | Using the resources of the University of Pittsburgh’s Brain Tissue Donation Program, 210 subjects were identified. Samples were obtained after consent from next-of-kin during autopsies conducted at the Allegheny County Medical Examiner’s Office (Pittsburgh, USA). The absence of lifetime psychiatric disorders was determined by an independent committee of experienced clinical research scientists using information from clinical records, toxicology results and a standardized psychological autopsy. Subjects of unwitnessed death were removed from the study. A total of 146 individuals with the following characteristics were analyzed in this study: mean (range) age of 50.7 (16-96) years, 78% male, 85% Caucasian, mean postmortem interval (PMI) for brain collection 17.3 (4.8-28) hours, mean pH 6.7 (5.8-7.6) and RNA integrity number (RIN) 8.0 (5.9-9.6). |
| 11. Human epidermal and dermal skin samples (population) | Kimball A. B., et al., 2018, J Am Acad Dermatol.  GSE112660, GSE139305 | One skin sample from forearm, cheek and buttock was taken from each of 154 female participants, aged between 20 and 74 years old. Samples were designed to collect during the working hours, between 9am to 5pm. LCM was performed to separate dermis from epidermis. There are 17 participants with one missing sample and one participant with two missing samples. |
| 12. Human GEO-LIHC (population) | GSE25097 | Profiles 268 HCC tumor, 243 adjacent non-tumor, 40 cirrhotic and 6 healthy liver samples. |
| 13. Human GEO-LUAD (population) | Selamat SA, et al., 2012, Genome Res.  GSE32863 | 58 lung adenocarcinoma and 58 adjacent non-tumor lung fresh frozen tissues were macrodissected, and total RNA was isolated to be analyzed using the Illumina HumanWG-6 v3.0 expression beadchip. |
| 14. Human GEO-KIRC (population) | Wozniak MB, et al., 2013, PLoS One.  GSE40435 | Conducte whole-genome expression profiling on 101 pairs of ccRCC tumours and adjacent non-tumour renal tissue from Czech patients using the Illumina HumanHT-12 v4 Expression BeadChips to explore the molecular variations underlying the biological and clinical heterogeneity of ccRCC. |
| 15. Human TCGA-BRCA, TCGA-THCA, TCGA-KIRC, TCGA-LIHC,  TCGA-LUAD,  TCGA-PRAD | http://firebrowse.org/ | The Cancer Genome Atlas (TCGA), a landmark cancer genomics program, molecularly characterized over 20,000 primary cancers and matched normal samples spanning 33 cancer types. This joint effort between NCI and the National Human Genome Research Institute began in 2006, bringing together researchers from diverse disciplines and multiple institutions. |
